# The chemorepellent, SLIT2, bolsters innate immunity against *Staphylococcus aureus*

**DOI:** 10.1101/2022.11.11.515814

**Authors:** Vikrant K. Bhosle, Chunxiang Sun, Sajedabanu Patel, Johannes Westman, Dustin A. Ammendolia, Noah Fine, Zhubing Li, Manraj Sharma, Judah Glogauer, Mariana I. Capurro, Nicola L. Jones, Michael Glogauer, Sergio Grinstein, Lisa A. Robinson

## Abstract

Neutrophils are essential for host defense against *Staphylococcus aureus* (*S. aureus*). The neuro-repellent, SLIT2, potently inhibits neutrophil chemotaxis, and might therefore be expected to impair antibacterial responses. We report here that, unexpectedly, neutrophils exposed to the N-terminal SLIT2 (N-SLIT2) fragment kill extracellular *S. aureus* more efficiently. N-SLIT2 amplifies reactive oxygen species production in response to the bacteria by activating p38 mitogen-activated protein kinase that in turn phosphorylates NCF1, an essential subunit of the NADPH oxidase complex. N-SLIT2 also enhances exocytosis of secondary granules. In a murine model of *S. aureus* skin and soft tissue infection (SSTI), local SLIT2 levels fall initially but increase subsequently, peaking ≈ 3 days after infection. Of note, neutralization of endogenous SLIT2 worsens SSTI. Temporal fluctuations in tissue SLIT2 levels may promote neutrophil recruitment and retention at the infection site and hasten bacterial clearance by augmenting neutrophil oxidative burst and degranulation. Collectively, these actions of SLIT2 coordinate innate immune responses to limit susceptibility to *S. aureus*.

**One-sentence summary:** SLIT2, a prototypic neuro-repellent, spatiotemporally coordinates host defense against *Staphylococcus aureus* infection.

## Introduction

*Staphylococcus aureus* (*S. aureus*) is a commensal bacterium as well as a skillful, facultative pathogen causing diverse human diseases ranging from localized skin and soft tissue infection (SSTI) to life-threatening disseminated sepsis (*1, 2*). Neutrophils, the most abundant subset of leukocytes in circulation, form a formidable first-line of defense against *S. aureus* invasion and spread (*3*). Accordingly, patients with neutropenia and defective neutrophil functions are highly susceptible to recurrent and more severe *S. aureus* infections (*4-6*). Infections caused by antibiotic-resistant strains of *S. aureus* have been steadily increasing worldwide, suggesting that strategies to enhance effective recruitment of neutrophils and their inherent bactericidal properties are needed to combat the morbidity and mortality associated with *S. aureus* infections (*7, 8*).

A recent transcriptomic study noted that mRNA encoding Slit guidance ligand 2 (SLIT2), a canonical neuro-repellent, is locally upregulated during *S. aureus*-induced mastitis (*9*). Among the 3 mammalian SLIT family members, SLIT1 is exclusively expressed in the nervous system (*10*); while SLIT2 and SLIT3 are also detected outside the nervous system (*11, 12*). We and others have previously reported that SLIT2 inhibits directed neutrophil migration *in vitro* as well as *in vivo* (*10, 13-16*). We also showed that the cleaved N-terminal fragment of SLIT2, N-SLIT2, acts via its receptor, Roundabout guidance receptor 1 (ROBO1), to attenuate inflammasome activation in macrophages by inhibiting macropinocytosis (*11*). Despite the inhibitory actions of N-SLIT2 on chemotaxis and macropinocytosis in innate immune cells, administration of N-SLIT2 *in vivo* does not confer increased susceptibility to bacterial infections (*15, 17*). It remains unknown if and how N-SLIT2 affects neutrophil responses during *S. aureus* infection.

Herein, we show that N-SLIT2 does not inhibit, but rather enhances, killing of extracellular *S. aureus* by neutrophils. In the presence of N-SLIT2, human and murine neutrophils respond more robustly to *S. aureus* by activating p38 mitogen-activated protein kinase (MAPK) signaling, enhancing production of extracellular reactive oxygen species (ROS) and release of secondary and tertiary granule contents by neutrophils. In a murine model of *S. aureus*-induced SSTI (*18*), we found that endogenous levels of SLIT2 protein decline significantly early on but rose to peak levels approximately 3 days after infection, and that blocking endogenous SLIT2-ROBO1 signaling at the site of infection enhances bacterial survival and worsens the infection. Our results suggest that changes in the levels of SLIT2 at local sites of infection may coordinate neutrophil recruitment, retention, and bactericidal responses to effectively and synergistically target *S. aureus*. Our work identifies SLIT2 as an endogenous regulator of neutrophil number and activity that coordinates immune responses vital to combat disseminated infection of bacterial pathogens.

## Results

### N-SLIT2 augments extracellular ROS production in response to *S. aureus*

We first investigated how SLIT2 affects neutrophil-mediated killing of *S. aureus*. Incubation of primary human neutrophils with bioactive N-SLIT2, but not inactive N-SLIT2ΔD2 which lacks ROBO1/2-binding D2 LRR domain (*19*), significantly reduced extracellular *S. aureus* as early as 30 minutes (Fig. 1A). Decreased extracellular bacteria could result from increased internalization via phagocytosis, and/or increased extracellular bacterial killing. Intriguingly, N-SLIT2 did not affect the ability of human neutrophils (Figs. 1B-D) or RAW264.7 murine macrophages to phagocytose *S. aureus* (Supplementary Figs. S1A-C). To determine whether N-SLIT2 decreased bacterial survival by stimulating phagocyte ROS production, a process dependent on NADPH Oxidase complex (NOX) activity, we incubated neutrophils with the pan-NOX-inhibitor, diphenyleneiodonium chloride (DPI). DPI partially restored extracellular bacterial survival in the presence of N-SLIT2 (Supplementary Fig. S1D). Next, we directly examined the effects of N-SLIT2 on extracellular ROS production by human neutrophils exposed to *S. aureus*. Neutrophils incubated with vehicle control or with N-SLIT2 alone had very low basal levels of ROS. As expected, *S. aureus* induced significant ROS production that was indistinguishable from that seen in cells incubated with *S. aureus* and bio-inactive N-SLIT2ΔD2 (Figs. 1E-F). Surprisingly, human neutrophils co-incubated with bio-active N-SLIT2 and *S. aureus* produced significantly more extracellular ROS than *S*.*aureus*-exposed neutrophils incubated with vehicle or N-SLIT2ΔD2 (Figs. 1E-F). We found that neutrophils exposed to another secondary ROS-inducing stimulus, namely phorbol-12-myristate-13-acetate (PMA), also produced more extracellular ROS in the presence of N-SLIT2 compared to N-SLIT2ΔD2 (Supplementary Fig. S1E). Similar observations were also noted for mouse bone marrow-derived neutrophils (BMDN) indicating that the effect of N-SLIT2 to further enhance extracellular ROS production by *S. aureus*-exposed neutrophils is not species-specific (Supplementary Fig. S1F). To determine whether the observed effects of N-SLIT2 occurred through the canonical ROBO1 receptor, we pre-incubated N-SLIT2 with the soluble N-terminal fragment of the ROBO1 receptor (N-ROBO1). As expected, N-ROBO1 fully blocked the ability of N-SLIT2 to boost extracellular ROS production in neutrophils exposed to *S. aureus* (Figs. 1G-H). Since activation of Rac GTPases is essential for optimal NOX function in neutrophils (*20, 21*), we next tested the effects of N-SLIT2 on Rac. N-SLIT2 alone failed to activate Rac but instead required a second stimulus, namely *S. aureus* (Supplementary Fig. S1G). Together, these results demonstrate that N-SLIT2 enhances neutrophil-mediated killing of *S. aureus*, partly by amplifying extracellular ROS production in a ROBO1-dependent manner.

**Figure 1.**
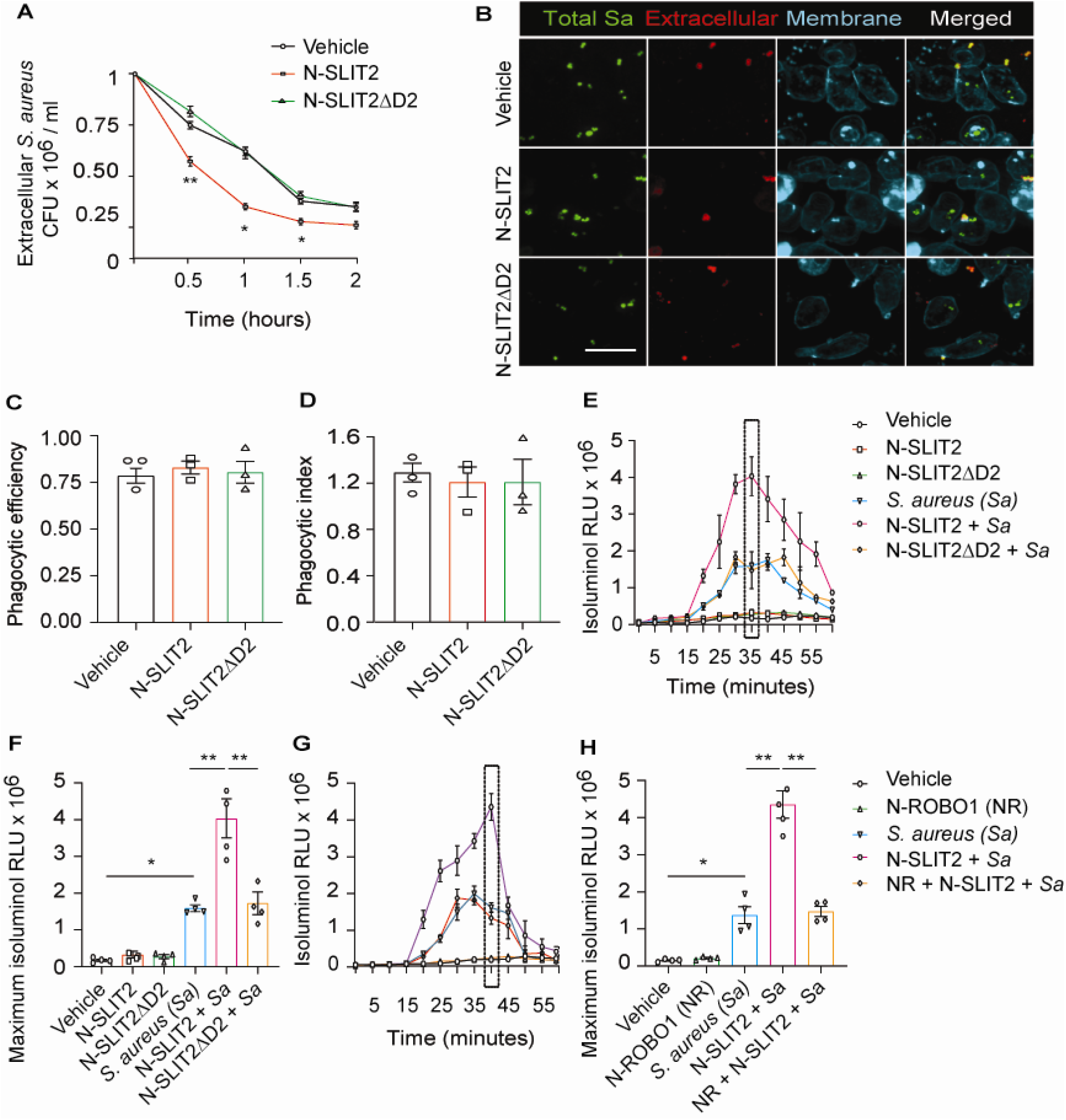
N-SLIT2 augments extracellular ROS production in response to *S. aureus*. (**A**) Neutrophils, isolated from healthy human donors, were incubated with vehicle (HBSS), N-SLIT2 (30 nM) or N-SLIT2ΔD2 (30 nM) for 15 min, followed by exposure to *S. aureus* (MOI 10) for the indicated times. Extracellular *S. aureus* counts were determined by serial dilution. n = 4. The statistical comparisons between N-SLIT2 and N-SLIT2ΔD2 groups are shown. p = 0.0072 (0.5 h), p = 0.0105 (1 h), p = 0.0478 (1.5 h), and p = 0.0852 (2 h). (**B**) Human neutrophils were treated with vehicle, N-SLIT2 or N-SLIT2ΔD2 for 15 min and then incubated with unoposonized *S. aureus* expressing GFP (MOI 10) for an additional 45 min. Extracellular bacteria were labeled using donkey anti-human IgG-Cy3. Neutrophil plasma membranes were labeled using Concanavalin A-AF647. At least 100 neutrophils per treatment were imaged. n = 3. The phagocytic efficiency (**C**) and index (**D**) were calculated. (**E**) The experiments were performed as in ‘A’ and extracellular ROS production was measured every 5 min using isoluminol relative luminescent units (RLU). n = 4. The averages of 4 experiments are shown. The timepoint with maximum extracellular ROS (35 min) is marked with a dotted rectangle. (**F**) Extracellular ROS production corresponding to maximum isoluminol RLU was compared among experimental groups. p = 0.0118 (vehicle vs *S. aureus*), p = 0.0099 (*S. aureus* vs N-SLIT2 + *S. aureus*), and p = 0.0055 (N-SLIT2 + *S. aureus* vs N-SLIT2ΔD2 + *S. aureus*). (**G**) Experiments were performed as described In (E) in parallel incubating N-SLIT2 (30 nM) with N-ROBO1 (NR; 90 nM) for 1 h at room temperature before adding to the cells. n = 4. Averages of all experiments are shown. The timepoint with maximum extracellular ROS (40 min) is marked with a dotted rectangle. (**H**) The timepoint with maximum isoluminol RLU was compared across experimental groups. p = 0.0250 (vehicle vs *S. aureus*), p = 0.0018 (*S. aureus* vs N-SLIT2 + *S. aureus*), and p = 0.0028 (N-SLIT2 + *S. aureus* vs N-ROBO1 + N-SLIT2 + *S. aureus*). Mean values ± SEM. * p < 0.05, and ** p < 0.01.

### N-SLIT2 primes NOX by p38-mediated phosphorylation of NCF1

We next studied how N-SLIT2 increases extracellular ROS production in neutrophils. We wondered whether N-SLIT2 induces NOX priming and extracellular ROS production by prompting phosphorylation and translocation of Neutrophil Cytosolic Factor 1 (NCF1; p47phox), a key component of the NOX complex, to the plasma membrane (*22, 23*). We found that N-SLIT2 increased phosphorylation of the conserved Ser^345^ residue of NCF1 in neutrophils (Figs. 2A-B), as well as in RAW264.7 macrophages (Supplementary Figs. S2A-B). As SLIT2-ROBO2 signaling was recently reported to activate NOX function in neurons by activating protein kinase C (PKC) (*24*), we examined the effects of N-SLIT2 on PKC activation in neutrophils. PKC activity in cells is tightly regulated by its phosphorylation (*25*). N-SLIT2 did not induce phosphorylation of PKC in neutrophils (Supplementary Figs. S2C-F). SLIT2 orthologues have been reported to activate p38 MAPK signaling in *Xenopus* and *C. elegans* neurons (*26, 27*). The p38 MAP kinases are also known to phosphorylate NCF1 at the Ser^345^ residue to mediate NOX priming (*28*). We, therefore, investigated the effects of N-SLIT2 on p38 MAPK signaling in mammalian phagocytes. N-SLIT2 potently induced p38 MAPK activation in both neutrophils (Figs. 2C-D) and macrophages (Supplementary Figs. S2G-H). Congruent with these results, incubation with two distinct pharmacologic inhibitors of p38 MAPKs blocked the observed N-SLIT2-induced increase in extracellular ROS in neutrophils exposed to *S. aureus* (Figs. 2E-F). These findings indicate that N-SLIT2 does not activate, but rather primes, the NOX complex in phagocytes to upregulate ROS production in response to injurious biologic and pharmacological secondary stimuli, including *S. aureus* and PMA, respectively.

**Figure 2.**
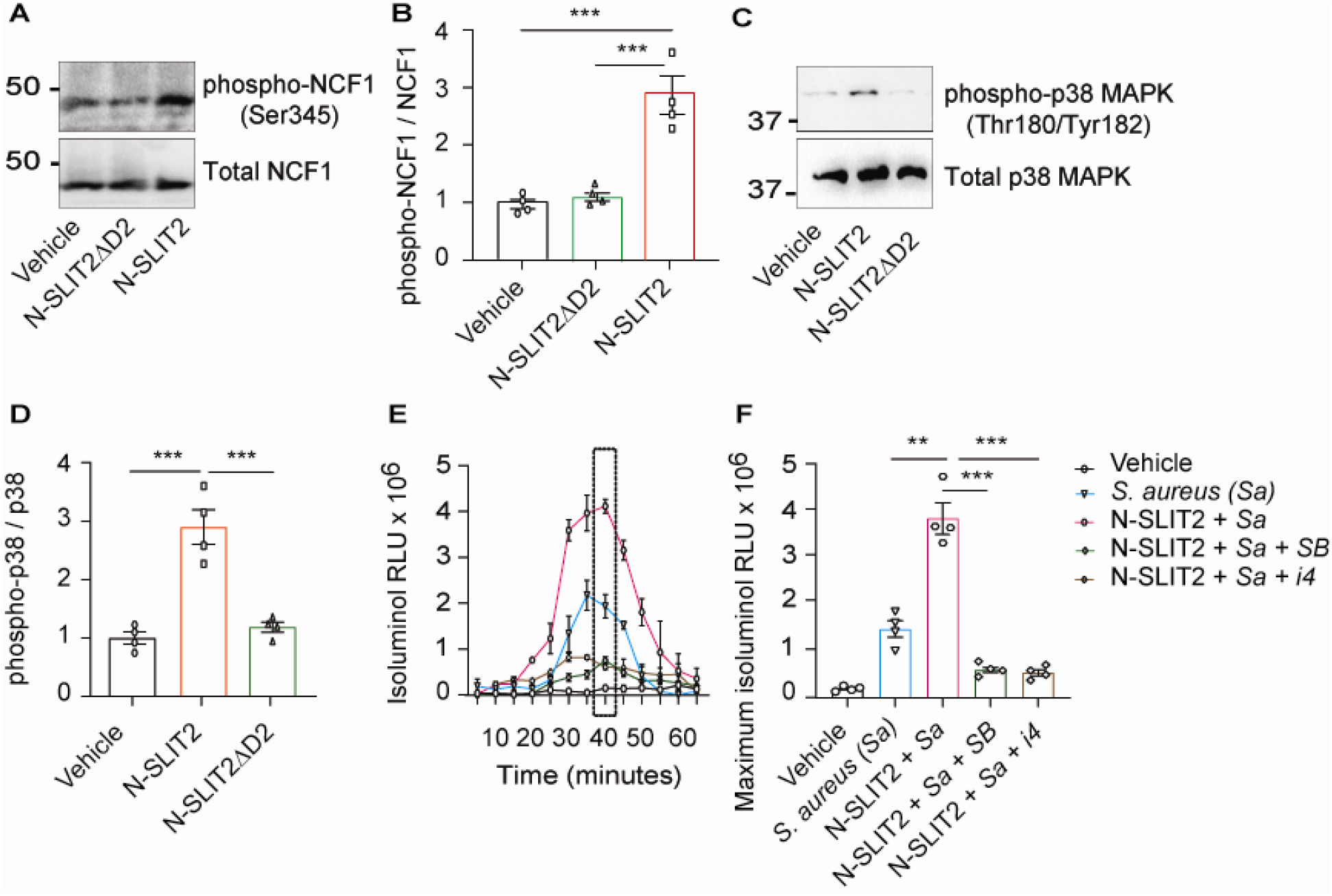
N-SLIT2 primes NOX by p38-mediated phosphorylation of NCF1. (**A**) Human neutrophils were exposed to vehicle, N-SLIT2 or N-SLIT2ΔD2 for 15 min and the protein lysates were immunoblotted for phospho-NCF1 (Ser345) and total NCF1 (Ser345). n = 4. A representative blot is shown. (**B**) Experiments were performed as in (A), densitometry performed, and the ratio of phospho-NCF1 / NCF1 obtained. n = 4. p = 0.0002 (vehicle vs N-SLIT2) and p = 0.0004 (N-SLIT2 vs N-SLIT2ΔD2). (**C**) Experiments were performed as in (A) and immunoblotting performed for phospho-p38 (Thr180/Tyr182) and total p38. n = 4. A representative blot is shown. (**D**) Experiments were performed as in (C), densitometry performed, and the ratio of phospho-p38 / p38 obtained. p = 0.0006 (vehicle vs N-SLIT2) and p = 0.0004 (N-SLIT2 vs N-SLIT2ΔD2). (**E**) Human neutrophils were incubated with vehicle, *S. aureus*, N-SLIT2 + *S. aureus*, N-SLIT2 + *S. aureu*s + SB203580 (SB, 10 μM), or N-SLIT2 + *S. aureus* + p38 MAPK Inhibitor IV (i4, 10 μM), and extracellular ROS were measured as described in Figure (1E). n = 4. The averages of 4 experiments are shown. The timepoint with maximum extracellular ROS (40 min) is marked with a dotted rectangle. (**F**) Experiments were performed as in (E) and extracellular ROS production at 40 min compared among groups. n = 4. p = 0.0036 (*S. aureus* vs N-SLIT2 + *S. aureus*), p = 0.0005 (N-SLIT2 + *S. aureus* vs N-SLIT2 + *S. aureus* + SB203580), and p = 0.0009 (N-SLIT2 + *S. aureus* vs N-SLIT2 + *S. aureus* + p38 MAPK Inhibitor IV). Mean values ± SEM. ** p < 0.01, and *** p < 0.001.

### N-SLIT2 enhances p38 MAPK-mediated exocytosis of secondary and tertiary granules

Since exposure to the NOX inhibitor, DPI, only partially reversed the effects of N-SLIT2 on extracellular *S. aureus* survival, we pondered whether other anti-microbial functions of neutrophils are also modified by N-SLIT2. In addition to the oxidative burst, degranulation has been shown to play an important role in bactericidal responses of neutrophils against *S. aureus* (*29, 30*). Neutrophils contain four types of granules, each containing different classes of anti-microbial peptides, of which secondary and tertiary granules are most important for eliminating *S. aureus* (*30-32*). To investigate the effects of N-SLIT2 on degranulation, we utilized a recently optimized flow cytometry protocol using specific cluster of differentiation (CD) markers of granule exocytosis (*33*). Another advantage of using a flow cytometry-based approach is that it circumvents the *in vitro* neutrophil isolation step which is known to affect cellular activation (neutrophil gating strategy; Fig. 3A). Neither *S. aureus* alone nor *S. aureus* together with N-SLIT2 induced primary granule (CD63) secretion (Supplementary Fig. S3A). Congruent with published literature (*34, 35*), the bacteria alone stimulated secondary granule (CD66b) exocytosis from neutrophils (Supplementary Fig. S3B). This effect was strikingly enhanced by exposure to N-SLIT2 (Fig. 3B and Supplementary Fig. S3B). Additionally, the actions of N-SLIT2 were completely obliterated by pharmacological inhibition of p38 MAPK signaling using SB 203580 or p38 MAPK Inhibitor IV, but not MEK1/2 signaling using PD 184161 (Fig. 3B). It is noteworthy that N-SLIT2 also augmented surface expression of *S. aureus*-induced CD18, which is stored in both secondary and tertiary granules, in a p38 MAPK dependent manner (Fig. 3C and Supplementary Fig. S3C) (*31, 32*). On the other hand, we found no differences in surface expression of CD16, which is stored in secretory vesicles, in any of the tested conditions, a finding in agreement with the priming-associated CD marker signature described for neutrophils in circulation and tissues (Supplementary Fig. S3D) (*33*). We next used surface expressions of CD66b and CD11b to calculate the percentage of primed neutrophils (Supplementary Figs. S3A and S3E) (*33*). Exposure of blood to N-SLIT2 and *S. aureus* together significantly increased the fraction of primed neutrophils as compared to *S. aureus* alone (Fig. 3D). Finally, we measured secretion of LL-37, a highly potent cationic anti-staphylococcal peptide that is selectively stored in neutrophil secondary granules in its pro-peptide form (*36, 37*). Incubation of human neutrophils with *S. aureus* resulted in release of LL-37, which was further amplified in the presence of N-SLIT2 and blocked by the p38 inhibitors (Fig. 3E). Collectively, our results elucidate that SLIT2-induced activation of p38 MAPK augments production of extracellular ROS, exocytosis of secondary granules, and secretion of the anti-bacterial LL-37 peptide from neutrophils in response to exposure to *S. aureus*.

**Figure 3.**
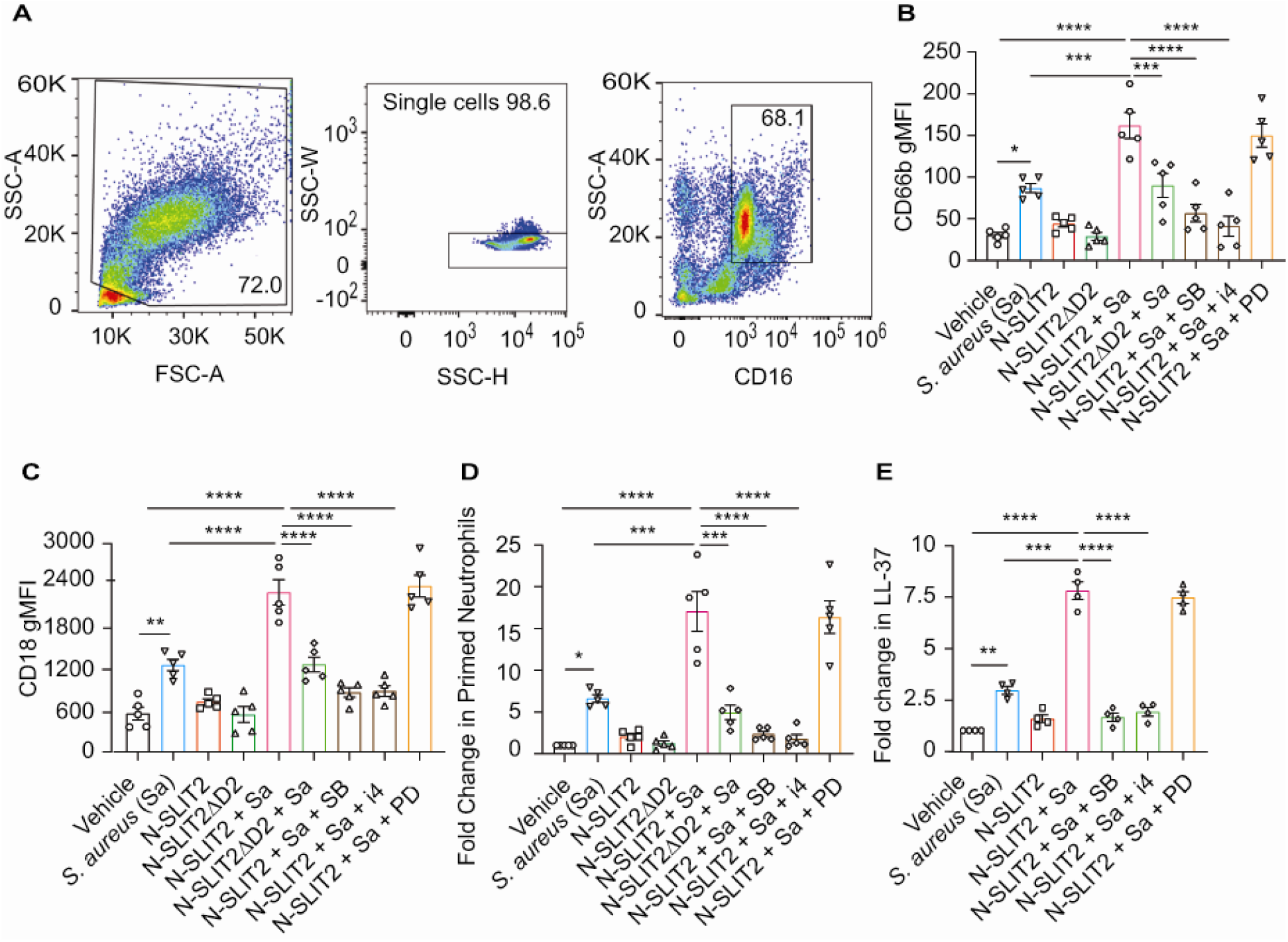
N-SLIT2 enhances p38 MAPK-mediated exocytosis of secondary and tertiary granules. (**A-D**) 100 μl whole blood from human subjects was exposed to different treatments for 15 min at 37°C, as indicated. The samples were immediately fixed on ice with 1.6% PFA and surface CD markers labeled. n = 5. (**A**) Gating strategy for human blood neutrophils: Red blood cells and dead cell debris were excluded based on FSC-A x SSC-A. Doublets were excluded based on SSC-A x SSC-W. Neutrophils were gated in whole blood leukocytes using CD16^high^ x SSC-A^high^. (**B**) Human neutrophils were exposed to vehicle, N-SLIT2, or N-SLIT2ΔD2 with or without the p38 MAPK inhibitors, SB 203580 (SB; 10 μM) or p38 MAPK Inhibitor IV (i4; 10 μM), or the MEK1/2 inhibitor PD 184161 (PD; 10 μM) for 15 min, followed by exposure to *S. aureus* (*Sa*) for another 15 min at 37°C, as indicated. Geometric mean fluorescence intensity (gMFI) for CD66b (secondary granules) is shown. p = 0.0122 (vehicle vs *Sa*), p < 0.0001 (vehicle vs N-SLIT2 + *Sa*) p = 0.0003 (*Sa* vs N-SLIT2 + *Sa*), p = 0.0006 (N-SLIT2 + *Sa* vs N-SLIT2ΔD2 + *Sa*), p < 0.0001 (N-SLIT2 + *Sa* vs N-SLIT2 + *Sa* + SB), and p < 0.0001 (N-SLIT2 + *Sa* vs N-SLIT2 + *Sa* + i4)002E (**C**) Neutrophils were treated as in (B) and gMFI for CD18 (secondary and tertiary granules) is noted. p = 0.0022 (vehicle vs *Sa*), p < 0.0001 (vehicle vs N-SLIT2 + *Sa*) p < 0.0001 (*Sa* vs N-SLIT2 + *Sa*), p < 0.0001 (N-SLIT2 + *Sa* vs N-SLIT2ΔD2 + *Sa*), p < 0.0001 (N-SLIT2 + *Sa* vs N-SLIT2 + *Sa* + SB), and p < 0.0001 (N-SLIT2 + *Sa* vs N-SLIT2 + *Sa* + i4). (**D**) Primed neutrophils were identified by cell surface labeling CD66b^high^ x CD11b^high^ and fold changes in % primed neutrophils relative to vehicle treatment are shown. p = 0.0246 (vehicle vs *Sa*), p < 0.0001 (vehicle vs N-SLIT2 + *Sa*) p = 0.0002 (*Sa* vs N-SLIT2 + *Sa*), p = 0.0008 (N-SLIT2 + *Sa* vs N-SLIT2ΔD2 + *Sa*), p < 0.0001 (N-SLIT2 + *Sa* vs N-SLIT2 + *Sa* + SB), and p < 0.0001 (N-SLIT2 + *Sa* vs N-SLIT2 + *Sa* + i4). (**E**) Human neutrophils were exposed to vehicle or N-SLIT2 with or without p38 MAPK inhibitors, SB or i4, or the MEK1/2 inhibitor PD for 15 min, then exposed to *S. aureus* (*Sa*) for another 15 min at 37°C, as indicated. Supernatants were collected and secreted LL-37 levels were measured using an ELISA. n = 4. p = 0.0092 (vehicle vs *Sa*), p < 0.0001 (vehicle vs N-SLIT2 + *Sa*) p = 0.0005 (*Sa* vs N-SLIT2 + *Sa*), p < 0.0001 (N-SLIT2 + *Sa* vs N-SLIT2 + *Sa* + SB), and p < 0.0001 (N-SLIT2 + *Sa* vs N-SLIT2 + *Sa* + i4). Mean values ± SEM. * p < 0.05, ** p < 0.01, *** p < 0.001 and **** p < 0.0001.

### Blocking endogenous SLIT2 exacerbates tissue injury in *S. aureus* SSTI

A recent transcriptomic screen has reported that *Slit2* mRNA is upregulated during *S. aureus*-induced mastitis (*9*). To determine whether this increase is conserved at a protein level during SSTI, we used an established murine model of *S. aureus* SSTI, which mimics community-acquired human infections (*18*). Surprisingly, we found that local SLIT2 levels were reduced as early as 12 hours after the infection (Fig. 4A). Following the initial decrease, the levels of SLIT2 gradually increased to reach a peak at 3 days (Fig. 4A). Unlike SLIT2, SLIT3 levels in the infected tissue remained significantly lower than in control mock-infected tissue for the first 2 days (Supplementary Fig. S4A). To ascertain the target of SLIT2 in *S. aureus* SSTI, we used soluble N-terminal fragment of human ROBO1 receptor (N-ROBO1) to block endogenous SLIT2 (*11, 38*). Based on the observed peak in endogenous SLIT2 levels at Day 3 after *S. aureus* infection, N-ROBO1 or control IgG was administered on Days 2 and 3 after infection (Fig. 4B). Strikingly, the bacterial counts were more than 2-fold higher in N-ROBO1-treated mice as compared to the IgG-treated counterparts (Fig. 4C). *S. aureus* infection in mice who received control IgG treatment or no other treatment resulted in microabscess formation with immune cell infiltration (Fig. 4D). In N-ROBO1-treated mice, *S. aureus* infection caused much more extensive tissue injury characterized by diffuse, rather than localized, inflammation (Figs. 4D-E and Supplementary Fig. S4B). Interestingly, in the absence of bacterial infection, neutralization of endogenous SLIT2 augmented immune cell infiltration in the skin but did not result in local tissue damage in the form of acanthosis (Fig. 4D). Our findings suggest that immediately after infection the rapid decrease in local levels of SLIT2 promotes infiltration of neutrophils into the site of infection. The later increase in SLIT2 levels may serve dual functions: local retention of neutrophils due to SLIT2’s chemorepellent actions, and direct effects on neutrophils to enhance their anti-staphylococcal killing responses.

**Figure 4.**
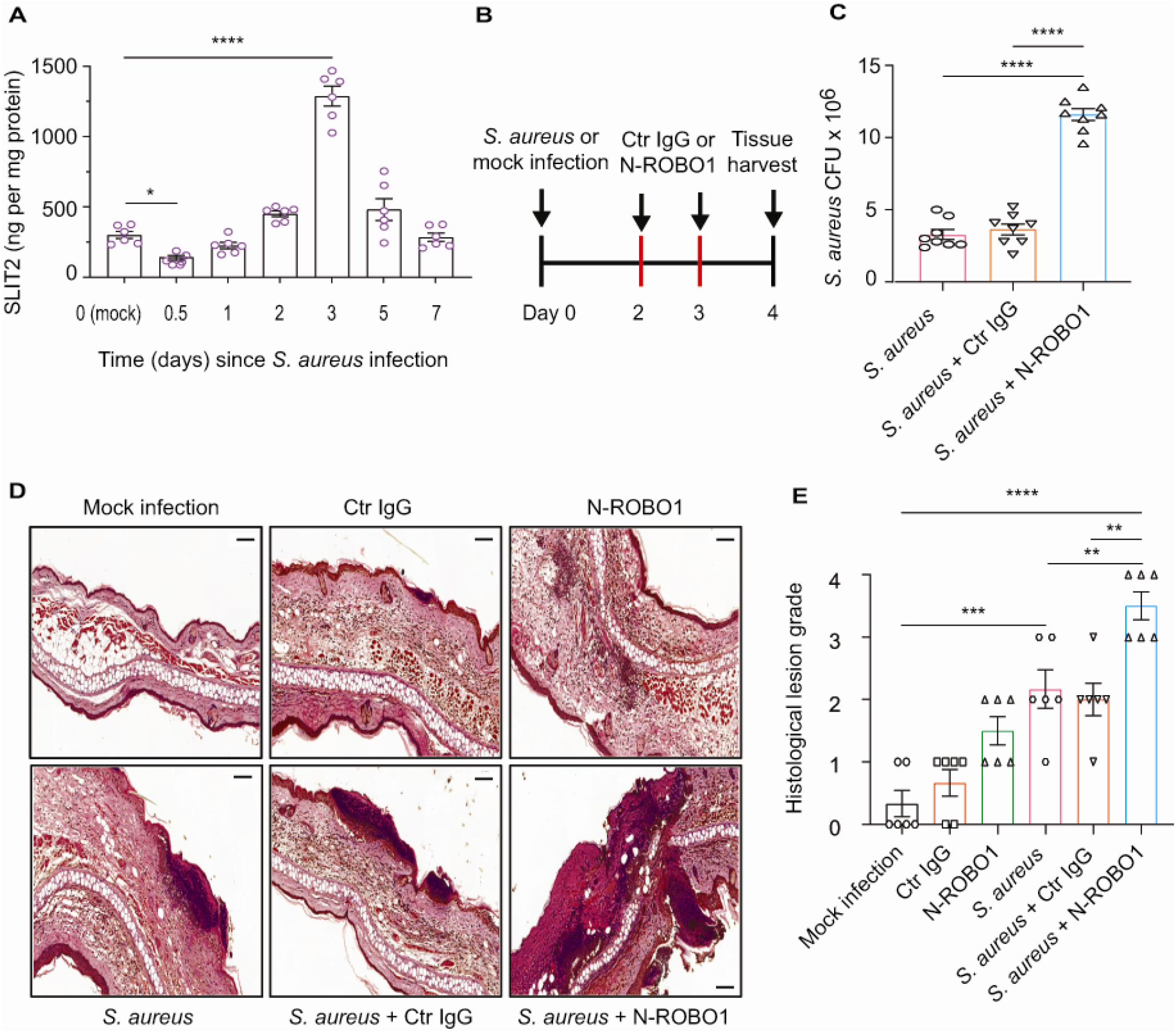
Blocking endogenous SLIT2 exacerbates tissue injury in *S. aureus* SSTI. (**A**) Ear skin samples were collected mock-infected (0) and *S. aureus*-infected mice at indicated time points in days, homogenized and tissue SLIT2 levels were measured using an ELISA. n = 6 mice per group. p = 0.0290 (Mock infection vs *S. aureus* 0.5 day), p = 0.002 (Mock infection vs *S. aureus* 3 days). (**B**) *In vivo* protocol to block endogenous SLIT2 during *S. aureus* SSTI. Ear pinnae were infected with 5 × 10^6^ CFU of *S. aureus* or mock-infected (saline injection) as indicated on day-0. On 2^nd^ and 3^rd^ day, local injections of either 7 μg of control (Ctr) IgG or N-ROBO1 were given at the site of infection. Skin samples were collected on 4^th^ day after the infection. (**C**) Samples were collected as described in (B) and were homogenized and *S. aureus* CFU were counted by serial dilution. p < 0.0001 (*S. aureus* vs *S. aureus* + N-ROBO1) and p < 0.0001 (*S. aureus* + Ctr IgG vs *S. aureu*s + N-ROBO1). N = 8 (**D-E**) Samples were collected as described in (B), fixed in formalin and stained with hematoxylin and eosin. Scale bar = 100 μm (**E**) Experiments were performed as in (D). The lesions were blindly scored on an ascending scale of severity (0-5). n = 6. p = 0.0002 (Mock infection vs *S. aureus*), p < 0.0001 (Mock infection vs *S. aureus* + N-ROBO1), p = 0.0060 (*S. aureus* vs *S. aureus* + N-ROBO1), p = 0.0016 (*S. aureus* + Ctr IgG vs *S. aureus* + N-ROBO1). Mean values ± SEM. * p < 0.05, ** p < 0.01, *** p < 0.001, and **** p < 0.0001.

## Discussion

We demonstrate here that N-SLIT2 augments extracellular ROS production by primary human and murine neutrophils in response to secondary stimulation with *S. aureus* and PMA. This effect is neutralized by incubating N-SLIT2 with N-ROBO1 and is therefore mediated via the canonical SLIT2-ROBO1 signaling pathway. These findings are in keeping with those of Wu *et al*. who reported that SLIT2 potentiates fMLP-induced oxidative burst in neutrophil-like HL-60 cells, without activating it on its own (*10*). Our work suggests that the inability of N-SLIT2 alone to induce ROS production by neutrophils could be explained by the lack of Rac activation (Supplementary Fig. S1G), as the latter is essential for activation of the NOX complex (*20*). We recently reported that SLIT2-ROBO1 signaling inhibits macropinocytosis in macrophages *in vitro* as well as *in vivo* (*11*). We show here that N-SLIT2 has no effect on phagocytosis of *S. aureus* by neutrophils and macrophages, in keeping with our previous observation that N-SLIT2 does not affect FcγR-mediated phagocytosis of opsonized particles by neutrophils (*15*). Together, these results suggest that N-SLIT2 can selectively promote some phagocyte functions while attenuating or not affecting others.

The phagocyte oxidative burst is a powerful tool for host defense against versatile pathogens such as *S. aureus* (*5, 6*). The oxidative burst can be further enhanced by priming agents, including tumor necrosis factor-α, which increase ROS production in response to secondary activating stimuli but have no effect on their own (*28*). In the resting state, three out of the five core components of the NOX complex, namely NCF1, NCF2, and NCF4, reside in the cytosol while the other two, CYBA and CYBB, are transmembrane proteins (*39*). We selectively investigated NCF1 (also known as p47phox) because of its well characterized role in promoting NOX activation at the plasma membrane, and therefore, extracellular ROS production (*23, 40*). The primed state of the NOX complex is characterized by phosphorylation of its cytosolic components, including NCF1, followed by their membrane translocation (*39*). We found that, upon exposure to N-SLIT2, NCF1 is phosphorylated at the conserved Serine^345^ residue, which in turn results in NOX priming (*28*) in phagocytes. To date, more than a dozen cellular protein kinases, including PKC and p38 MAPKs, have been implicated in NCF1 phosphorylation (*22*). Terzi *et al*. postulated that neuronal SLIT2-ROBO2 signaling could activate PKC (*24*). We observed that in neutrophils, N-SLIT2 did not activate PKC. These findings could be explained by the observation that in a cell-free system, the PKC-interacting protein AKAP79 binds to C-termini of ROBO2/3, but fails to bind to the C-terminus of ROBO1 (*41*). Accordingly, we and others previously reported that primary human and murine neutrophils express ROBO1 (*13, 42*), but not ROBO2 (*42*). Our studies revealed that N-SLIT2 induced p38 activation in human neutrophils and that inhibition of p38 signaling potently attenuated the observed N-SLIT2-mediated increase in oxidative burst. These results are in accordance with the observation that SLIT2 and its orthologues can stimulate p38 MAPK signaling in non-mammalian neurons (*26, 27*). Our results are also in agreement with a recent study reporting hyperactivation of p38 MAPK in mice genetically overexpressing SLIT2 (*43*). Interestingly, during the oxidative burst, PKC and p38 MAPK target distinct serine residues in the NCF1 C-terminus for phosphorylation (*28, 44, 45*). This might explain how N-SLIT2 (which activates p38) and PMA (which activates PKC) act synergistically during ROS production by human neutrophils. Together, these findings show that N-SLIT2-ROBO1 signaling primes the phagocyte NOX complex via p38 MAPK-mediated phosphorylation of NCF1.

An ever growing body of evidence reveals phenotypic and functional heterogeneity of neutrophils in the circulation as well as in tissues (*33, 46, 47*). As some of the neutrophil subtypes are sensitive to *in vitro* isolation methods, a more representative snapshot of their inherent properties is enabled by optimized flow cytometry protocols that use whole blood preparations (*33, 48*). We found that in addition to priming neutrophils, N-SLIT2 also enhanced exocytosis of secondary granules after exposure to *S. aureus*, stimulating release of LL-37, an antimicrobial cathelicidin peptide which is highly effective in killing *S. aureus* extracellularly at nanomolar concentrations (*36*). Our results are congruent with previous reports that exposure to *S. aureus* increases CD66b surface levels in neutrophils (*49*), but we noted that this effect was further augmented by concurrent exposure to both N-SLIT2 and *S. aureus*. This can be explained by our observation that N-SLIT2 activates p38 MAPK and primes the NOX complex. In neutrophils, priming of the NOX complex by p38 MAPK results in exocytosis of secondary granules, and LL-37 is stored in its pro-peptide form in secondary granules (*37, 50, 51*). Indeed, we observed that inhibition of NOX partially blocked the effects of N-SLIT2, while inhibition of p38 MAPK abolished both extracellular ROS production as well as secondary granule release.

Although N-SLIT2 enhanced *S. aureus*-induced exocytosis of secondary granules in neutrophils, it did not alter cell-surface levels of the phagocytic receptor, CD16. Similarly, exposure to N-SLIT2 did not change cell-surface levels of CD63, a protein stored in primary (azurophilic) granules of neutrophils (*31*). Our results are in line with studies showing that neither *S. aureus* nor p38 activation induce primary granule release from neutrophils (*34, 51*). Together, these findings show that N-SLIT2-induced activation p38 MAPK is responsible for secretion of secondary granules which contain the anti-staphylococcal peptide, LL-37 (Supplementary Fig. S4C).

Using a murine model of community-associated *S. aureus*-induced SSTI, we found that endogenous levels of SLIT2 protein at the site of infection were significantly reduced at 0.5 day after infection, and subsequently increased to reach a peak at 3 days post-infection. Local levels of SLIT3 were also reduced at 0.5 day but did not increase more than those in uninfected controls. Congruent with our findings, local reduction in *SLIT3* mRNA levels was recently reported in patients after *S. aureus*-induced SSTI (*52*). Importantly, we found that locally blocking endogenous SLIT2 at time points when it is most abundantly expressed resulted in enhanced *S. aureus*-induced SSTI. These results are consistent with our previous observation that neutralization of endogenous SLIT2 enhances macrophage macropinocytosis and inflammatory cytokine production *in vivo* in a peritonitis model (*11*). Taken together, these results indicate that host-derived SLIT2 plays an important role in the spatiotemporal modulation of innate immune responses during bacterial infection. The rapid initial decrease in SLIT2 (and SLIT3) at the site of infection may serve to augment the immediate recruitment of neutrophils to combat the infection, while the rise in SLIT2 several days later may promote localized retention of recruited neutrophils and enhancement of their bactericidal properties. In this manner, spatiotemporal regulation of SLIT2/ROBO1 activity may potently direct bacterial killing and confer effective host protection against pathogenic infection. Further studies are needed to explore the precise mechanisms by which levels of SLIT2 are modulated in the face of bacterial infection and to explore the potential of SLIT2 as a novel anti-microbial therapeutic. Given mounting concerns about the growing resistance of bacterial pathogens to conventional pharmacologic antibiotic treatment, SLIT2 may represent an attractive alternative or synergistic therapeutic strategy.

## Materials and Methods

All antibodies used in this study are listed in Supplementary Table 1. All chemicals, reagents, and assay kits are listed in Supplementary Table 2.

### Cell Culture

RAW264.7 (TIB-71™) cells were purchased from American Type Culture Collection (ATCC; Manassas, VA, USA). Cells were cultured in Dulbecco’s Modified Eagle’s Medium (DMEM; Wisent, St-Bruno, QC, Canada) supplemented with 10% heat-inactivated fetal bovine serum (FBS; Wisent) with 1% antibiotic/antimycotic (Wisent) at 37°C and 5% CO_2_. HEK293F cells were grown in FreeStyle 293 Expression Medium (GIBCO™, Thermo Fisher Scientific, Mississauga, ON, Canada). All cell lines tested negative for mycoplasma using the Mycoplasma Plus™ PCR kit (#302008, Agilent Technologies, Santa Clara, CA, USA).

### Production of recombinant SLIT2 proteins

Large-scale production of bio-inactive human N-SLIT2ΔD2 was performed by transfecting HEK293F cells with human N-SLIT2ΔD2 cDNA (1 ug/mL) using PEI reagent (Polyethyleneimine, linear, M.W. 25,000, Thermo Fisher Scientific) (*19*). After 5 days, culture medium was loaded onto HisPur Ni-NTA resin (Thermo Fisher Scientific), washed with imidazole (35 mM), eluted with imidazole (250 mM), and desalted using a Pierce Zeba™ desalting column (7K MWCO; Thermo Fisher Scientific). Protein’s molecular weight was determined by immunoblotting using anti-6XHIS-HRP conjugated antibody. Protein activity was assayed by using spreading assay in RAW264.7 cells, as described previously (*11*). Recombinant human N-SLIT2 was purchased from PeproTech (Cranbury, NJ, USA). Endotoxin levels in all preparations were measured using ToxinSensor™ Chromogenic LAL Endotoxin Assay Kit (GenScript, Piscataway, NJ, USA) and were less than 0.05 EU/ml.

### Human neutrophil isolation

All reagents were filtered through Detoxi-Gel™ Endotoxin Removing Gel Columns (Thermo Fisher Scientific) prior to neutrophil isolation. Human neutrophils were isolated using a density-gradient separation method (*53*). Briefly, 30 ml of blood was collected in sterile BD Vacutainer^®^ EDTA (BD Biosciences, Mississauga, ON, Canada) tubes and layered on the top of PolymorphPrep™ (Progen, Wayne, PA, USA) at a 1:1 ratio. The layered mixture was spun at 500 g (acceleration 1 and deceleration 0) for 30 min to separate polymorphonuclear neutrophils (PMN) from mononuclear cells. PMN were washed with Hank’s balanced salt solution with calcium and magnesium (HBSS+/+; Wisent) and then subjected to red blood cell lysis using 0.2% NaCl followed by 1.6% NaCl for 30 s each. Neutrophils were resuspended in HBSS+/+ and used within 1 h for all experiments.

### Murine BMDN isolation

C57BL/6 mice aged 8 to 12 weeks were purchased from Charles River Laboratories (St-Constant, QC, Canada). Mice were euthanized by CO_2_ inhalation, and their tibias and femurs were resected and cut at both ends. Bone marrow cells were extracted and added to Percoll (Sigma-Aldrich, Oakville, ON, Canada) gradient solutions of 82%/65%/55% as previously described (*13*). The solution was centrifuged, and murine neutrophils were collected from 82%/65% interface. Red blood cell lysis was performed as described above.

### *S. aureus* killing assays

*Staphylococcus aureus subsp. Aureus Rosenbach (S. aureus*) (ATCC 25923™) was purchased from ATCC. The extracellular bactericidal activity of neutrophils was assayed as described previously (*54*), with some modifications. Freshly isolated neutrophils (1 × 10^5^) were incubated with vehicle (HBSS+/+), N-SLIT2, or N-SLIT2ΔD2 for 15 min at 37°C and then incubated with unoposonized *S. aureus* (MOI 10) for an additional 15 min at 37°C. End-over rotation (6 rpm) was used to prevent neutrophil clumping. Tubes were centrifuged at 100 g for 5 min and neutrophil pellets were discarded. Supernatant samples containing the extracellular bacteria were collected. Bacterial colony forming units (CFU) were determined using serial dilution followed by overnight culture in Tryptic Soy Agar (TSA) with 5% sheep blood (Remel, Lenexa, KS, USA) (*54*).

### *S. aureu*s phagocytosis assays

The GFP-expressing strain *of S. aureus* USA300 was grown at 37°C overnight with shaking in Todd-Hewitt broth (BD Biosciences) with erythromycin (3 μg/mL) for selection (*55*). Just before the phagocytosis assay, RAW264.7 macrophages or freshly-isolated human neutrophils were incubated with vehicle, N-SLIT2 or N-SLIT2ΔD2 for 15 min at 37°C.

Overnight *S. aureus* USA300 were diluted to OD_600nm_ 0.05 in Todd-Hewitt broth containing erythromycin and cultivated until logarithmic growth (OD_600nm_ 0.5) at 37°C. Bacterial aggregates were dispersed using a 27-gauge needle (BD Biosciences) prior to infection. Unopsonized *S. aureus* were centrifuged onto RAW264.7 macrophages (MOI 10) or human neutrophils (MOI 10), and phagocytosis was allowed to proceed for 45 min at 37°C under 5% CO_2_. Samples were washed three times in sterile PBS and fixed in 3% paraformaldehyde (PFA) (Electron Microscopy Sciences, Hatfield, PA, USA) for 15 min at room temperature (RT). Samples were blocked with block buffer, consisting of PBS containing 2% FBS and 2% bovine serum albumin (BSA) for 30 min at RT. To label bound but non-internalized bacteria, samples were incubated with human total IgG for 30 min at room temperature. Non-internalized bacteria were visualized using donkey anti-human IgG-Cy3 for 30 min at room temperature. To label the cell nuclei, samples were permeabilized with Triton-X100 (0.2%) for 10 min and incubated with DAPI for 10 min. The surface of RAW264.7 cells was labeled with Acti-stain-AF670 for 10 min at RT. The surface of neutrophils was labeled with Concanavalin A-AF647 for 30 min.

Confocal images were acquired using a spinning disk system (WaveFX; Quorum Technologies Inc., Puslinch, ON, Canada). The instrument consists of a microscope (Axiovert 200M; Zeiss, Toronto, ON, Canada), scanning unit (CSU10; Yokogawa Electric Corp, Calgary, AB, Canada), electron-multiplied charge-coupled device (C9100-13; Hamamatsu Photonics), five-line (405-, 443-, 491-, 561-, and 655-nm) laser module (Spectral Applied Research, Richmond Hill, ON, Canada), and filter wheel (MAC5000; Ludl) and is operated by Volocity software version 6.3 (PerkinElmer, Waltham, MA, USA). Confocal images were acquired using a 633/1.4-N.A. oil objective (Zeiss) coupled to an additional 1.53 magnifying lens and the appropriate emission filter. Phagocytic Index and Phagocytic Efficiency were quantified as described below (*56*).

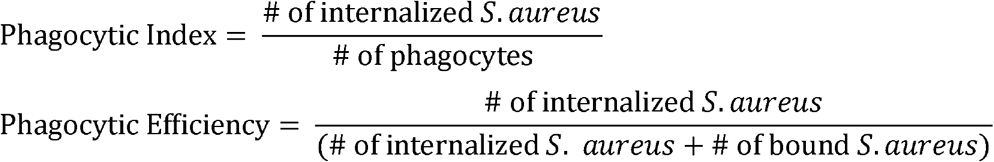

### Extracellular ROS production

Extracellular ROS levels were measured using an isoluminol-based chemiluminescence assay (*57*). Human or murine neutrophils (5 x10^5^) were incubated with isoluminol (3 mmol/L) and horseradish peroxidase (6 U) in a 96-well plate and exposed to N-SLIT2 or N-SLIT2ΔD2 for 15 min immediately followed by a secondary stimulus consisting of *S. aureus* (MOI 10) or PMA (100 nM) at 37°C. Chemiluminescence was measured at every 5 minutes (for a total of 45 minutes) using a Varioskan LUX Plate Reader (Thermo Fisher Scientific).

### Rac G-LISA

Activation of Rac1/2/3 was measured by G-LISA as previously described (*11*). Following the experimental treatment, neutrophils were lysed using boiling (95°C) lysis buffer with protease inhibitors, and assays were performed according to the manufacturer’s instructions (*11*). Readings were obtained at 490□nm using a Plate Reader.

### Immunoblotting

Neutrophils were lysed in boiling RIPA lysis buffer (60 mM Tris, pH 7.5, 10% Glycerol, 1% TX-100, 150 mM NaCl, 0.5% Na-deoxycholate, 0.1% SDS, 1mM EDTA) and 1X protease and phosphatase inhibitor. Protein lysate (30 μg) with 2X Laemmli buffer (in 1:1 ratio) was loaded onto a 4%–20% gradient gel (Bio-Rad, Mississauga, ON, Canada). The gel was transferred onto a PVDF membrane using the Trans-Blot Turbo Transfer System (Bio-Rad). Membranes were blocked for 1 hour at RT in blocking buffer (5% BSA in PBS-T), with a solution containing primary antibodies. After washing and addition of the secondary antibody, membranes were developed using SuperSignal™ West Dura photographed using the Bio-Rad GelDoc system with Image Lab software. The denistometric quantifications were performed using ImageJ software, version 1.51v (*58*).

### Flow Cytometry

All treatments were performed with whole blood for 15 min at 37°C. Immediately following the treatment, the samples were fixed with PFA (1.6%) for 15 min on ice. RBCs were lysed three times with Pharm Lyse™ (BD Biosciences) for 10 minutes on ice. Cells were resuspended in flow-assisted cell sorting (FACS) buffer (HBSS-/-, 1% BSA, 2 mM EDTA), labeled with an antibody cocktail for 30 min on ice in the dark, and washed 3 times with FACS buffer. Sample acquisition was performed using a Sony Spectral Cell Analyzer SA3800 in a standardization mode. At least 1 × 10^5^ gated events were acquired per sample (*33, 48*). Data were analyzed using FlowJo software (BD Biosciences, version 10). Human multicolor flow cytometry panel and gating were done as previously described (*33*) using anti-human CD66b antibody, a marker of secondary granules.

### Epicutaneous *S. aureus* infection model

The epicutaneous model of *S. aureus* infection has been described previously (*18*). C57BL/6 mice aged 8 to 12 weeks were anesthetized using an intraperitoneal injection of ketamine (50 mg/kg) and xylazine (5 mg/kg). The left ear was cleaned with 70% ethanol and pricked 10 times with an *S. aureus*-coated or sterile uncoated (mock infection) 25G BD SafetyGlide™ needle (#305901, BD Biosciences). The mean inoculation dose of *S. aureus* was determined to be 5 × 10^6^ CFU/lesion, as determined by plating serial dilutions of homogenates from ear pinnae 4 h after the infection. Mice were monitored twice daily. For N-ROBO1 *in vivo* experiments, in some experiments, after anesthesia mice received subcutaneous ear injections of N-ROBO1 or control IgG (7□μg) on days 2 and 3 after inoculation with *S. aureus*. Animals were euthanized using CO_2_ inhalation on day 4 after infection; ear pinnae were cleaned, resected in 500 μl of sterile PBS, and homogenized using a Fisherbrand™ 150 Handheld Homogenizer (#15-340-167, Thermo Fisher Scientific). Bacterial CFUs were determined by serial dilution.

### ELISA

Following treatment of neutrophils supernatants were collected and snap-frozen in liquid nitrogen. ELISA to measure LL-37 was performed as per the manufacturer’s protocol. A microplate reader was used to read absorbance at 450nm. In some experiments, murine ear pinnae were homogenized and levels of SLIT2 and SLIT3 measured by ELISA as per the manufacturers’ protocols (*11*). Optical density readings were measured on a microplate reader at 450nm (VersaMax 190, Molecular Devices).

### Histopathology

5 μm thick coronal sections (3 sections per sample) were embedded in Paraffin. Two sections per sample were counterstained with hematoxylin and eosin (H&E) at The Centre for Phenogenomics (Toronto, ON, Canada). The H&E Slides were scanned using 3DHistech Slide Scanner (The Hospital for Sick Children Imaging Facility) at 40x magnification and examined under blinded conditions on a scale of 1 to 5 as defined previously (*18*). The scoring system is indicated by:

0, no evidence of inflammation
1, minimal to mild focal dermal inflammatory infiltrates with or without dermal acanthosis;
2, mild multifocal dermal infiltrates with or without intraepidermal microabscesses;
3, multifocal to diffuse moderate dermal inflammatory infiltrates extending to the skeletal muscles with or without intraepidermal microabscesses;
4, marked dermal inflammatory infiltrates extending to the skeletal muscle and cartilage with or without intraepidermal microabscesses;
5, full thickness auricular necrosis or massive inflammation with myositis and chondritis.

### Study approvals

The protocol (#1000070040) for human blood collection from healthy adult donors was reviewed and approved by The Hospital for Sick Children (SickKids, Toronto, ON, Canada) Research Ethics Board. Written, informed consent was obtained from all participants prior to their participation. The animal studies were reviewed and approved by SickKids Animal Care Committee (ACC) (protocol #1000049735 – *S. aureus* infection) and The Centre for Phenogenomics (Toronto, ON, Canada) ACC (protocol #21-0326 – murine neutrophil isolation).

### Statistics

GraphPad Prism 9 was used for all statistical analyses (San Diego, CA, USA). All data are presented as mean +/- standard error of mean (SEM). Shapiro-Wilk test was first used to confirm a normal (Gaussian) distribution. For normal distribution, one-way ANOVA followed by Tukey’s post hoc analysis was used. Data with non-Gaussian distribution were analyzed using a Mann-Whitney test. Statistical tests are described in the Fig. legends. A *p*-value of less than 0.05 was considered statistically significant. The *p*-values are reported up to 4 decimal places.

## Acknowledgments

We thank Dr. Ronald S. Flannagan (University of Western Ontario, London, ON, Canada) for S. aureus (USA300) strain expressing GFP (*55*). We thank The Centre for Phenogenomics for histopathology services. We thank Dr. Spencer A. Freeman (The Hospital for Sick Children, Toronto, ON, Canada) for helpful scientific discussions.

## Author contributions

Conceptualization: VKB, SG, LAR

Methodology: VKB, CS, JW, NF, MIC, MG

Investigation: VKB, CS, SP, JW, NF, ZL, MS, DAA, JG

Resources: LAR, SG, NLJ, MG

Supervision: LAR

Writing - original draft: VKB

Writing - review & editing: VKB, SG, LAR

All authors reviewed and approved the final submission.

## Data and materials availability

All data are available in the main text or the supplementary materials. The requests for materials should be addressed to the corresponding author (LAR).

