## Supplementary Figures S1-4 and Tables 1-2 for "The chemorepellent, SLIT2, bolsters innate immunity against *Staphylococcus aureus*"

Vikrant K. Bhosle *et al.*

**This PDF file includes:**

Supplementary Figs. S1 to S4
Supplementary Tables S1 to S2

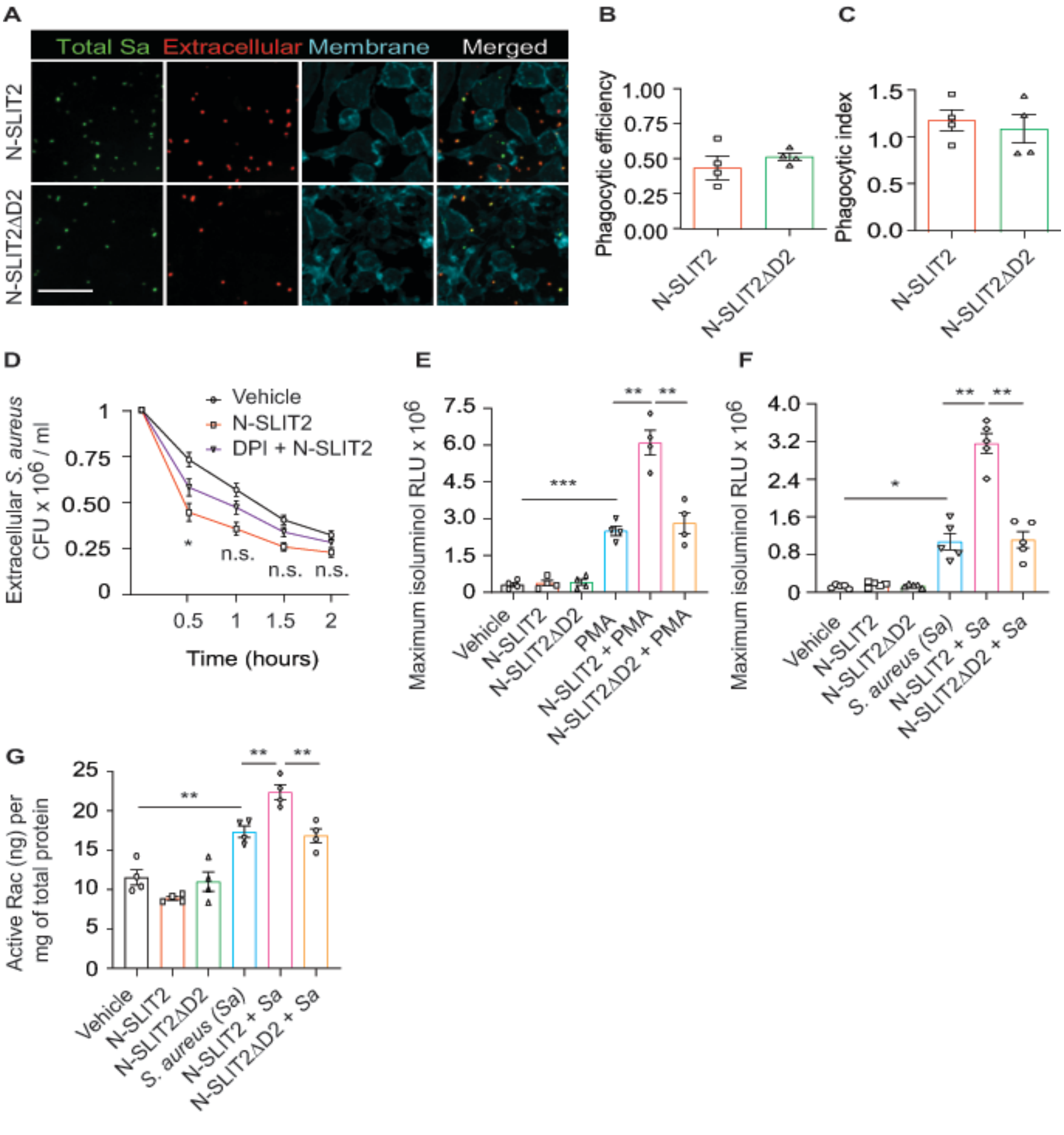


**Fig. S1**

(**A**) RAW264.7 cells were incubated with vehicle, N-SLIT2 (30 nM) or N-SLIT2ΔD2 (30 nM) for 15 min and then with unoposonized *S. aureus* expressing GFP (MOI 10) for an additional 45 min. Extracellular bacteria were labeled using donkey anti-human IgG-Cy3. Neutrophil plasma membranes were stained using Acti-stain-AF670. At least 100 neutrophils per experimental condition were imaged. n = 3. The phagocytic efficiency (**B**) and index (**C**) were calculated. (**D**) Experiments were performed as in Figure (1A). In some experiments, cells were pre-incubated with DPI (10 μM) for 5 min and then incubated with N-SLIT2 (30 nM) in the presence of DPI for an additional 15 min. n = 4. Statistical comparisons between N-SLIT2 and DPI + N-SLIT2 groups are p = 0.0172 (0.5 h), p = 0.0525 (1 h), p = 0.2775 (1.5 h), and p = 0.6207 (2 h). (**E**) Experiments were performed as in Figure (1E) using PMA (200 nM) as a secondary stimulus instead of *S. aureus*. Maximum isoluminol RLU were compared. n = 4. p = 0.0006 (vehicle vs PMA), p = 0.0065 (PMA vs N-SLIT2 + PMA), and p = 0.0092 (N-SLIT2 + PMA vs N-SLIT2ΔD2 + PMA). (**F**) Murine bone marrow-derived neutrophils were treated as described in Figure (1E). n = 4. p = 0.0158 (vehicle vs *S. aureus*), p = 0.0064 (*S. aureus* vs N-SLIT2 + *S. aureus*), and p = 0.0082 (N-SLIT2 + *S. aureus* vs N-SLIT2ΔD2 + *S. aureus*). (**G**) Human neutrophils were treated as described in (F) and active levels of Rac were measured using a Rac1/2/3 calorimetric G-LISA. n = 4. p = 0.0024 (vehicle vs *S. aureus*), p = 0.0088 (*S. aureus* vs N-SLIT2 + *S. aureus*), and p = 0.0038 (N-SLIT2 + *S. aureus* vs N-SLIT2ΔD2 + *S. aureus*). Mean values ± SEM. * p < 0.05; ** p < 0.01, *** p < 0.001, and n.s. not significant.

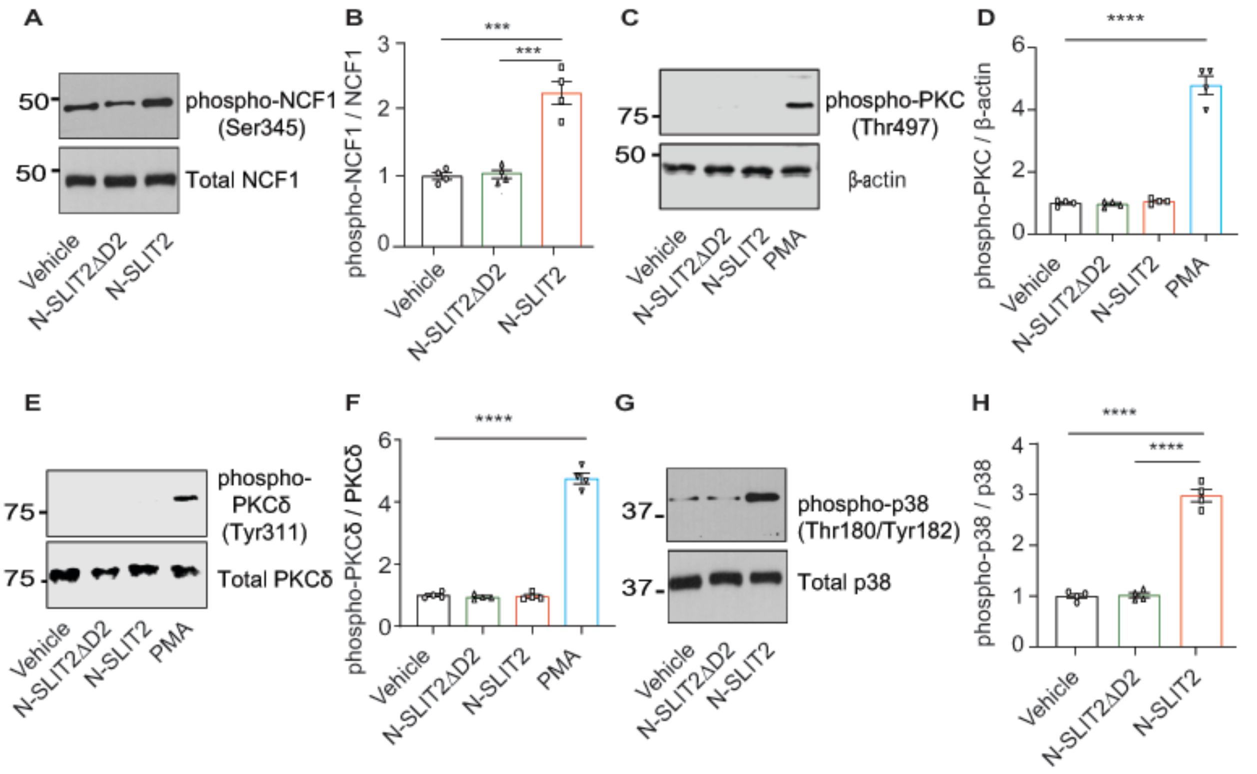


**Fig. S2**

(**A**) RAW264.7 cells were incubated with vehicle, N-SLIT2 or N-SLIT2ΔD2 for 15 min, and the protein lysates were immunoblotted for phospho-NCF1 (Ser345) and total NCF1 (Ser345). n = 4. A representative blot is shown. (**B**) Experiments were performed as in (A), densitometry performed, and the ratio of phospho-NCF1/NCF1 obtained. p = 0.0007 (vehicle vs N-SLIT2) and p = 0.0008 (N-SLIT2 vs N-SLIT2ΔD2). (**C**) Representative immunoblot showing lysates from human neutrophils treated with vehicle, N-SLIT2, N-SLIT2ΔD2, or PMA (200 nM) for 15 min and probed with phospho-PKC (Thr497) and β-actin. n = 4. (**D**) Densitometry analysis of phospho-PKC/β-actin ratios for experiments in (C). $p < 0.0001$ (vehicle vs PMA). (**E**) Representative immunoblot showing lysates from human neutrophils treated with vehicle, N-SLIT2, N-SLIT2ΔD2, or PMA (200 nM) for 15 min and probed with phospho-PKCδ (Tyr311) and Total PKCδ. n = 4. (**F**) Densitometry analysis of phospho-PKCδ/PKCδ ratios for experiments in (E). $p < 0.0001$ (vehicle vs PMA). (**G**) RAW264.7 cells were treated as described in (A) and the protein lysates were immunoblotted for phospho-p38 (Thr180/Tyr182) and total p38. n = 4. A representative blot is shown. (**H**) Densitometry analysis of phospho-p38/ p38 ratios for experiments in (E). $p < 0.0001$ (vehicle vs N-SLIT2) and $p < 0.0001$ (N-SLIT2 vs N-SLIT2ΔD2). *** $p < 0.001$ and **** $p < 0.0001$.

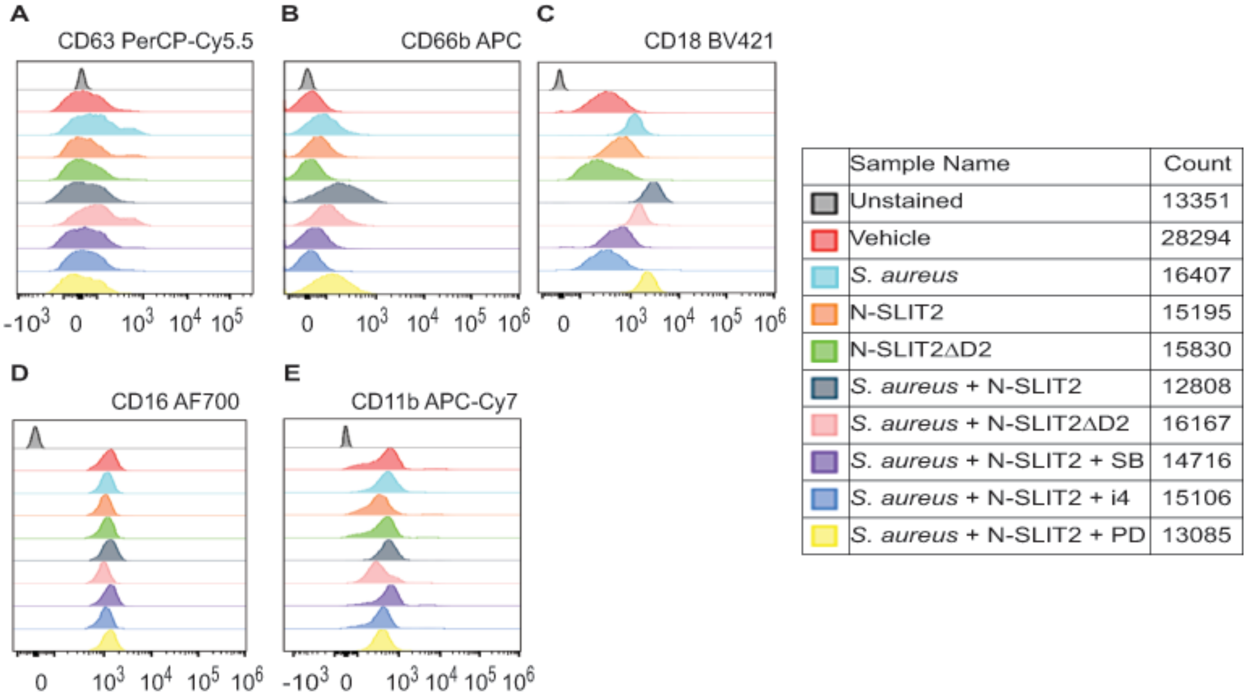


**Fig. S3**

Representative histograms are shown for human blood neutrophils with different treatments as described in Figure 3 to detect cell-surface expression of the following CD markers: (**A**) CD63 (**B**) CD66b (**C**) CD18 (**D**) CD16 and (**E**) CD11b.

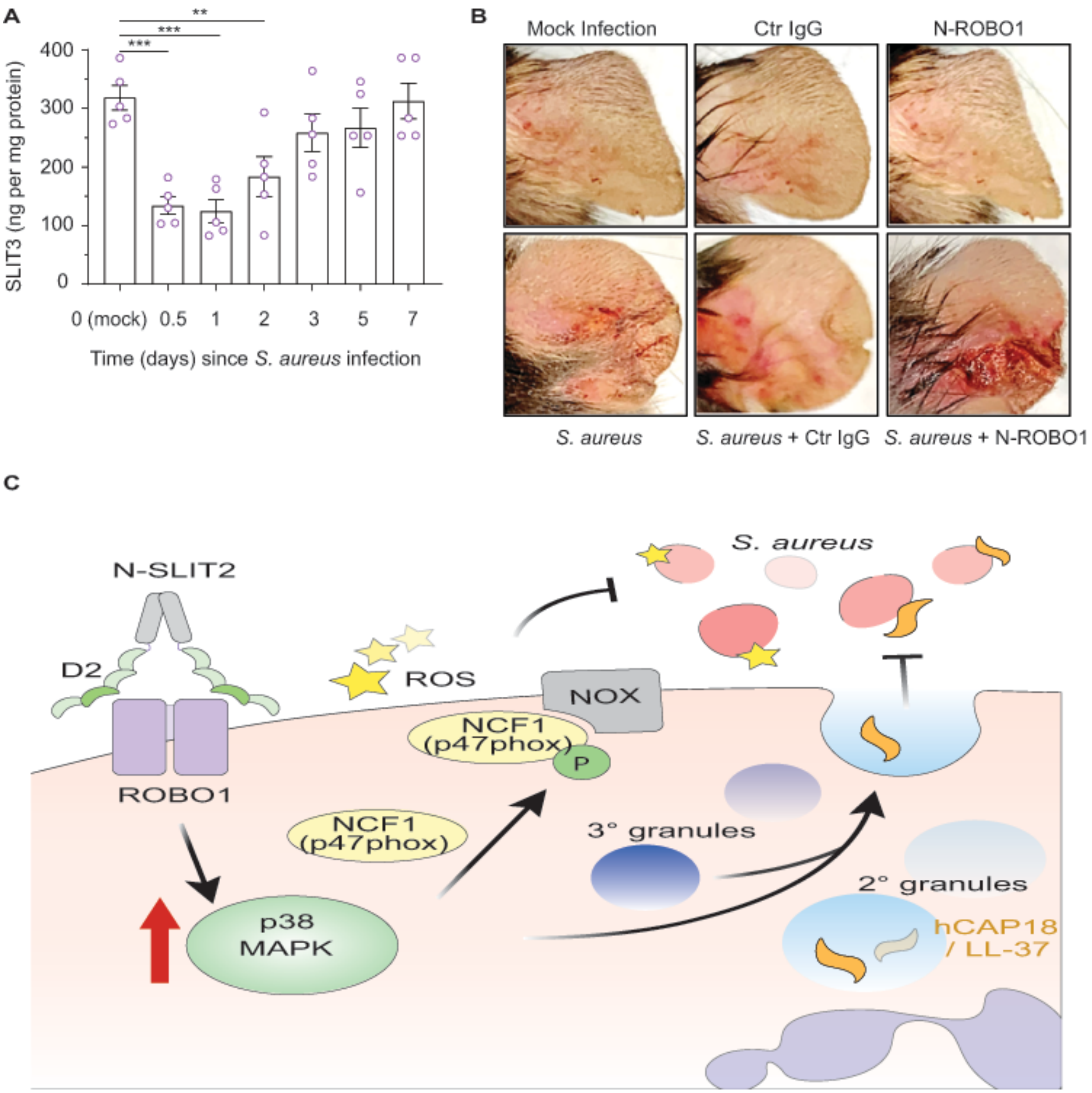


**Fig. S4**

(A) Ear skin samples were collected as described in (Figure 4A) and tissue SLIT3 levels were measured using an ELISA. n = 5. p = 0.0002 (0 vs *S. aureus* 0.5 day), p = 0.0003 (0 vs *S. aureus* 1 day), and p = 0.0067 (0 vs *S. aureus* 2 days). (B) Representative images of gross pathology of ear tissue from animals treated as described in (Figure 4B). ** $p < 0.01$, and *** $p < 0.001$. (C) Proposed mechanism of N-SLIT2's anti-bacterial action: The binding of Leucine-rich repeat D2 domain of N-SLIT2 to cell-surface ROBO1 results in activation of p38 MAPK signaling in neutrophils. Active p38 MAPK phosphorylates cytosolic NCF1 (p47phox), which is in its resting state, inducing translocation to the plasma membrane, thereby converting NCF1 to its primed state, forming a multi-protein NOX complex. In the presence of secondary stimuli such as *S. aureus* (and PMA), N-SLIT2-induced phosphorylation of NCF1 results in increased extracellular oxidative burst by neutrophils. The activation of p38 MAPK also augments exocytosis of secondary and tertiary granules. Secondary granules contain hCAP-18 which is cleaved extracellularly into its active form, the anti-microbial peptide, LL-37. Together, the N-SLIT2-mediated upsurge in both ROS production and LL-37 secretion promote enhanced extracellular killing of *S. aureus*.

**Table S1 Antibodies used in the study**

| REAGENT OR RESOURCE | SOURCE | IDENTIFIER (CATALOG # and RRID) | APPLICATION NOTES |
|---|---|---|---|
| **Primary Antibodies** | | | |
| Mouse Anti β-Actin Clone AC-15 | Sigma-Aldrich, Oakville, ON, Canada | #A5441 AB_476744 | Western blot (WB)- 1:2000 1 hr @ room temperature (RT) |
| Rabbit Anti Phospho-p38 MAPK (Thr180/Tyr182) | Cell Signaling Technology, Danvers, MA, USA | #9211 AB_331641 | WB- 1:2000 1 hr @ RT |
| Rabbit Anti Total p38 MAPK | Cell Signaling Technology, Danvers, MA, USA | #9212 AB_330713 | WB- 1:2000 Overnight (O/N) @ 4°C |
| Rabbit Anti Phospho-p47phox (NCF1) (Ser345) | Thermo Fisher Scientific, Mississauga, ON, Canada | #PA5-37806 AB_2554414 | WB- 1:1000 O/N @ 4°C |
| Rabbit Anti p47phox (NCF1) Clone G.207.2 | Thermo Fisher Scientific, Mississauga, ON, Canada | #MA5-14778 AB_10989232 | WB- 1:1000 O/N @ 4°C |
| Rabbit Anti Phospho-PKC Pan (Thr497) | Thermo Fisher Scientific, Mississauga, ON, Canada | #PA5-38418 AB_2555019 | WB- 1:1000 1 hr @ RT |
| Rabbit Anti Phospho-PKCδ (Tyr311) | Cell Signaling Technology, Danvers, MA, USA | #2055 AB_330876 | WB- 1:2000 1 hr @ RT |
| Rabbit Anti PKCδ | Cell Signaling Technology, Danvers, MA, USA | #2058 AB_10694655 | WB- 1:2000 O/N @ 4°C |
| Mouse HRP-conjugated Anti-His tag Clone AD1.1.10 | R&D Systems, Inc. Minneapolis, MN, USA | #MAB050H AB_357354 | WB- 1:2000 1 hr @ RT |
| Human Total IgG | Sigma-Aldrich, Oakville, ON, Canada | #I4506 AB_1163606 | Phagocytosis- 1:1000 in block buffer, 30 min RT |
| Donkey Anti-human IgG-Cy3 | Jackson ImmunoResearch Labs, West Grove, PA, USA | #709-165-149 AB_2340535 | Phagocytosis- 1:1000 in block buffer, 30 min RT |
| InVivoMAb human $IgG_1$ Isotype Control | Bio X Cell, Lebanon, NH, USA | #BE0297 AB_2687817 | *In vivo* - 7 µg per injection per mouse |

| | | | |
|---|---|---|---|
| **Secondary Antibodies** | | | |
| Peroxidase-conjugated AffiniPure Goat Anti-Mouse IgG (H+L) | Jackson ImmunoResearch Labs, West Grove, PA, USA | #115-035-003 AB_10015289 | WB- 1:10000 1 hr @ RT |
| Peroxidase-conjugated AffiniPure Goat Anti-Rabbit IgG (H+L) | Jackson ImmunoResearch Labs, West Grove, PA, USA | #111-035-144 AB_2307391 | WB- 1:10000 1 hr @ RT |
| **Flow Cytometry Antibodies** | | | |
| Normal Mouse IgG | Sigma-Aldrich, Oakville, ON, Canada | #12-371 AB_145840 | Flow cytometry (FC)- 2 µg per sample, To block 20 min @ 4°C |
| PE anti-human CD16 Clone 3G8 | BioLegend, San Diego, CA, USA | #980102 AB_2616616 | FC- 1 µl per 50 µl final volume, 30 min @ 4°C in the dark |
| APC/Cyanine7 anti-human CD11b Clone ICRF44 | BioLegend, San Diego, CA, USA | #301342 AB_2563395 | FC- 1.25 µl per 50 µl final volume, 30 min @ 4°C in the dark |
| BV421 anti-Human CD18 Clone 6.7 | BD Biosciences, Mississauga, ON, Canada | #562871 AB_2737855 | FC- 1.25 µl per 50 µl final volume, 30 min @ 4°C in the dark |
| PerCP/Cyanine5.5 anti-human CD63 Clone H5C6 | BioLegend, San Diego, CA, USA | #353020 AB_2561685 | FC- 1.25 µl per 50 µl final volume, 30 min @ 4°C in the dark |
| Pacific Blue™ anti-human CD14 Clone HCD14 | BioLegend, San Diego, CA, USA | #325616 AB_830689 | FC- 2.5 µl per 50 µl final volume, 30 min @ 4°C in the dark |
| APC anti-human CD66b, eBioscience™ Clone G10F5 | Thermo Fisher Scientific, Mississauga, ON, Canada | #17-0666-42 AB_2573152 | FC- 1.25 µl per 50 µl final volume, 30 min @ 4°C in the dark |

**Table S2 Chemicals, Reagents, Assays (ELISA) and Cell lines**

| RESOURCE | SOURCE | IDENTIFIER (CATALOG #) | Application |
|---|---|---|---|
| **Reagents and Chemicals** | | | |
| 2x Laemmli Sample Buffer | Bio-Rad Laboratories, Mississauga, ON, Canada | 1610737 | Immunoblotting |
| 4′,6-diamidino-2-phenylindole dihydrochloride (DAPI) | Thermo Fisher Scientific, Mississauga, ON, Canada | D1306 | Immunostaining (nuclear stain) |
| Acti-stain-AF670 | Universal Biologicals, Cambridge, UK | PHDN1-A | Phagocytosis |
| Antibiotic-Antimycotic solution (100X) | Wisent Inc., St-Bruno, QC, Canada | 450-115-EL | Cell culture |
| BD Pharm Lyse™ lysing solution (10X) | BD Biosciences, Mississauga, ON, Canada | 555899 | Flow Cytometry |
| BD Vacutainer® EDTA Tubes | BD Biosciences, Mississauga, ON, Canada | 366643 | Blood collection |
| BluElf prestained protein ladder | FroggoBio, Concord, ON, Canada | PM008 | Immunoblotting |
| Bovine Serum Albumin (BSA) | Sigma-Aldrich, Oakville, ON, Canada | A4503 | Blocking (phospho-protein Immunoblotting) |
| Concanavalin A-AF647 | Thermo Fisher Scientific, Mississauga, ON, Canada | C21421 | Phagocytosis |
| Detoxi-Gel™ Endotoxin Removing Gel | Thermo Fisher Scientific, Mississauga, ON, Canada | 20339 | Endotoxin removal |
| Detoxi-Gel™ Endotoxin Removing Gel Columns | Thermo Fisher Scientific, Mississauga, ON, Canada | 20344 | Endotoxin removal |
| Dimethyl sulfoxide (DMSO)<br>CAS #67-68-5 | Sigma-Aldrich, Oakville, ON, Canada | D8418 | Organic solvent |
| Diphenyleneiodonium | Sigma-Aldrich, | D2926 | NADPH Oxidase |

| | | | |
|---|---|---|---|
| chloride (DPI) CAS # 4673-26-1 | Oakville, ON, Canada | | Inhibitor |
| Dulbecco's Modified Eagle's Medium (DMEM) | Wisent Inc., St-Bruno, QC, Canada | 319-007-CL | RAW264.7 cell culture |
| Ethylenediaminetetraacetic acid (EDTA), 0.5M, pH 8.0, Sterile CAS #60-00-4 | Bio-World, Dublin, OH, USA | 40520000 | Flow Cytometry buffer ingredient |
| Fetal bovine serum (FBS) Premium Quality, heat inactivated, Endotoxin <1, Hemoglobin <25 | Wisent Inc., St-Bruno, QC, Canada | 080-150 | Cell culture |
| Gibco™ FreeStyle™ F17 Expression Medium | Thermo Fisher Scientific, Mississauga, ON, Canada | A1383501 | HEK293F cell culture |
| Formalin solution, neutral buffered, 10% | Sigma-Aldrich, Oakville, ON, Canada | HT501128 | Histology |
| Hank's Buffered Salt Solution with calcium and magnesium (HBSS +/+) | Wisent Inc., St-Bruno, QC, Canada | 311-513-CL | Buffered solution |
| Isoluminol (4-Aminophthalhydrazide) CAS #3682-14-2 | Sigma-Aldrich, Oakville, ON, Canada | A8264 | Extracellular ROS measurement |
| Narketan, ketamine hydrochloride injection | CDMV, Saint-Hyacinthe, QC, Canada | Not applicable | Anesthesia |
| Rompun® (xylazine injection) | CDMV, Saint-Hyacinthe, QC, Canada | Not applicable | Anesthesia |
| MS-SAFE™ | Sigma-Aldrich, Oakville, ON, Canada | MSSAFE | Protease and Phosphatase Inhibitor cocktail |
| p38 MAPK Inhibitor IV CAS #1638-41-1 | Cayman Chemical, Ann Arbor, MI, USA | 22219 | p38 MAPK inhibitor |
| Paraformaldehyde 16% solution CAS #50-00-0 | Electron Microscopy Sciences, Hatfield, PA, USA | 15710 | Fixative |
| PD 184161 CAS #212631-67-9 | Cayman Chemical, Ann Arbor, MI, USA | 10012431 | MEK1/2 inhibitor |
| Percoll® | Sigma-Aldrich, Oakville, ON, Canada | P1644 | Murine neutrophil isolation |
| Phorbol 12-myristate 13-acetate (PMA) | Sigma-Aldrich, Oakville, ON, | P8139 | Neutrophil activation |

| | | | |
|---|---|---|---|
| CAS #16561-29-8 | Canada | | |
| Phosphate Buffered Saline with calcium and magnesium (PBS +/+) | Wisent Inc., St-Bruno, QC, Canada | 311-011-CL | Buffered solution |
| Phosphate Buffered Saline without calcium and magnesium (PBS -/-) | Wisent Inc., St-Bruno, QC, Canada | 311-010-CL | Buffered solution |
| Poly-D-lysine<br>CAS #27964-99-4 | Sigma-Aldrich, Oakville, ON, Canada | P6407 | Coverslip coating |
| PolymorphPrep™ | Progen, Wayne, PA, USA | 1895 | Human neutrophil isolation |
| PVDF Membranes | Thermo Fisher Scientific, Mississauga, ON, Canada | 88518 | Immunoblotting |
| Rat serum | Sigma-Aldrich, Oakville, ON, Canada | R9759 | Flow (blocking solution ingredient) |
| Recombinant Human N-SLIT2 | PeproTech, Cranbury, NJ, USA | 150-11 | Neutrophil treatment |
| Recombinant Human ROBO1 Fc Chimera (N-ROBO1) | R&D Systems, Minneapolis, MN, USA | 8975-RB | *In vivo* |
| Restore™ PLUS Western Blot Stripping Buffer | Thermo Fisher Scientific, Mississauga, ON, Canada | 46430 | Immunoblotting |
| SB 203580<br>CAS #152121-47-6 | Sigma-Aldrich, Oakville, ON, Canada | S8307 | p38 MAPK inhibitor |
| Sodium Chloride Injection USP (0.9%) | Pfizer Canada Inc., Kirkland, QC, Canada | 4888010 | *In vivo* |
| Sodium Chloride salt | Sigma-Aldrich, Oakville, ON, Canada | S9888 | RBC lysis during neutrophil isolation |
| Triton X-100 | Bio-Rad Laboratories, Mississauga, ON, Canada | 1610407 | RIPA lysis buffer |
| Tryptic Soy Agar (TSA) with 5% sheep blood | Remel, Lenexa, KS, USA | R01200 | Agar plate for *S. aureus* |
| Tryptic Soy Broth (TSB) | Sigma-Aldrich, Oakville, ON, Canada | 22092 | *S. aureus* liquid growth medium |

| | | | |
|---|---|---|---|
| TWEEN® 20 Detergent CAS 9005-64-5 | Sigma-Aldrich, Oakville, Canada | 655204 | Immunoblotting |
| Zeba™ Spin Desalting Column | Thermo Fisher Scientific, Mississauga, ON, Canada | 89893 | Protein purification |
| **Assay kits including ELISA** | | | |
| Bio-Rad Protein Assay Kit | Bio-Rad Laboratories, Mississauga, ON, Canada | 5000001 | Immunoblotting |
| G-LISA® Rac Activation Assay | Cytoskeleton, Inc., Denver, CO, USA | BK125 | Rac1/2/3 activation assay |
| Human LL-37 ELISA | Hycult Biotech, Uden, Netherlands | HK321 | ELISA |
| Mouse SLIT2 ELISA Kit | Cusabio, Wuhan, P.R. China | CSB-E11039m | ELISA |
| SLIT3 ELISA kit | Lifespan Biosciences Seattle, WA, USA | LS-F-7173 | ELISA |
| ToxinSensor™ Chromogenic LAL Endotoxin Assay Kit | GenScript, Piscataway, NJ, USA | L00350 | Endotoxin |
| **Cell lines** | | | |
| FreeStyle™ 293-F Cells (HEK293F) | Thermo Fisher Scientific, Mississauga, ON, Canada | R79007 | SLIT2 purification |
| RAW264.7 | American Type Culture Collection (ATCC), Manassas, VA, USA | TIB-71™ | Murine macrophage cell line |
| **Bacterial strains** | | | |
| *Staphylococcus aureus* GFP (PMID: 30619165) | Dr. Ronald S. Flannagan (University of Western Ontario, London, ON, Canada) | USA300 strain | Phagocytosis |
| *Staphylococcus aureus subsp. Aureus Rosenbach* | American Type Culture Collection (ATCC), Manassas, VA, USA | ATCC® 25923™ strain | *S. aureus* (all experiments except phagocytosis) |
